# Population structure reduces the benefits from partner choice in mutualism

**DOI:** 10.1101/068445

**Authors:** Erol Akçay

## Abstract

Mutualisms are key drivers of evolutionary and ecological processes. Understanding how different species can evolve to interact in mutually beneficial ways is an important goal of evolutionary theory, especially when the benefits require costly investments by the partners. For such costly investments to evolve, some sort of fitness feedback mechanism must exist that more than recoups the direct costs. Several such feedback mechanisms have been explored both theoretically and empirically, yet we know relatively little how they might act together, as they probably do in nature. In this paper, I model the joint action of three of the main mechanisms that can maintain symbiotic cooperation: partner choice by hosts, population structure amongst symbionts, and undirected rewards from hosts to symbionts. My results show that population structure reduces the benefit from partner choice to hosts. It may help or hinder beneficial symbionts and create positive or negative frequency dependence depending on the nature of host rewards to the symbiont. Strong population structure also makes it less likely that host choosiness and symbiont cooperation will be jointly maintained in a population. The intuition behind my results is that all else being equal, population structure reduces local variation available to the host to choose from. Thus, population structure is not always beneficial for the evolution of cooperation between species. My results also underscore the need to do full analyses of multiple mechanisms of social evolution to uncover their interactions to uncover the interactions between them.

## 1 Introduction

Mutualisms are crucial mediators of important ecological and evolutionary processes, including ecosystem functions [1], disease dynamics in the wild and in humans [2], biodiversity [3, 4], and responses of ecosystems to changing environments [5]. Mutualistic interactions can arise simply as byproducts of adaptations in one species benefiting others [6], but many do involve traits that impose a direct cost on their bearers and directly benefit only other species. Such traits create a trade-off or conflict between the partners, because increasing a partner’s benefit (all else being equal) means decreasing one’s own [7–11]. This conflict is in tension with the opportunity for mutual benefit, and a large body of theoretical [reviewed in 8, 9, 12] and empirical work [e.g., 13–22] focuses on elucidating mechanisms that can resolve this tension and enable the evolutionary maintenance of mutual benefit in nature.

In general, mutualisms with costly investments can be evolutionarily stable if some mechanism creates a fitness feedback to a focal individual from investments that directly benefit the partner [12]. The most straightforward type of feedback is the direct coupling of partners’ fitnesses by vertical transmission of symbionts [23]. In the absence of such coupling, several other mechanisms exist that can generate fitness feedbacks favoring co-operative investments. These mechanisms include non-random association between co-operative partners through population structure [24] or through active partner choice [8, 25], and phenotypically plastic investments (termed phenotypic feedbacks [12]) that respond to benefits provided by a partner to create a return to cooperative investments. Phenotypic feedbacks can be specifically directed towards more cooperative partners [26,27], or undirected, accruing to all partners, but according to the total benefit received by the focal individual. Each of these mechanisms of fitness feedback have been extensively studied theoretically and empirically, but largely in isolation from each other, or under the assumption that they act additively [28]. As a result, we know relatively little about how multiple mechanisms of fitness feedbacks interact with each other, and whether such interactions should change our understanding of mutualism evolution.

Past theoretical work indicates that interactions between multiple types of feedbacks can be decisive for whether cooperation between species can evolve. Akçay and Simms [29] showed in the legume-rhizobium mutualism that both directed feedbacks (as produced by a negotiation process) and undirected ones are needed to maintain cooperation in rhizobia. Similarly, Van Cleve and Akçay [30] showed that phenotypic feedbacks and population structure within and between species reinforce each other’s effect, consistent with results from within-species social evolution theory [7, 31, 32]. Here, I focus on three major types of fitness feedback: partner choice, population structure, and different kinds of phenotypic feedbacks from the host.

Partner choice, i.e., hosts preferentially associating and rewarding more beneficial symbionts [8, 25], has long been recognized as a mechanism that can select for cooperation. Empirical work further shows that there is usually variation in host expression of partner choice [e.g. 16, 33, 34], and recent work has started to quantify more precisely the variation across [21, 35] and within species [36]. Recent theoretical work has also suggested that discriminating genotypes of hosts might coexist with non-discriminating ones [37, 38], which can explain how symbiont variability is maintained within populations. The latter is both an empirically observed phenomenon in need of an explanation [39], and a theoretical requirement for the maintenance of costly partner choice mechanisms [40].

In parallel, theory shows that when host reward their symbionts according to the benefits received, population structure amongst symbionts can favor cooperative investments because cooperative symbionts benefit from feedbacks induced by other cooperative neighbors that they tend to co-occur with [24, 41, 42]. However, the effects of partner choice in structured populations remains underexplored, except for a couple of recent studies by Bever et al. [43] and Verbruggen et al. [17] that show the role of within-host population structure in facilitating or hindering host plant partner choice of mycorrhizae. No previous study has formally considered how population-scale genetic structure of symbionts affects the evolutionary consequences of partner choice combined with phenotypic feedbacks.

Intuitively, a host can only exercise partner choice amongst symbionts available to it locally, and therefore effective partner choice requires local variation within the symbiont population. However, for a given overall level of genetic variation, stronger genetic structure (e.g., due to limited dispersal) will diminish local genetic variation. Therefore, we might expect partner choice in structured populations to be less effective than in well-mixed populations. On the other hand, as mentioned above, population structure can favor cooperation through kin-selected effects.

To explore how these potentially conflicting effects play out, I develop a general model of a choosy host interacting with a patch structured population of symbionts. My results show that population structure does reduce the benefit to the host from partner choice. On the symbiont side, the effect of population structure on the selection for symbiont co-operation depends on the nature of the rewards from the host. When symbionts receive a fixed reward from the host per infection, population structure disfavors the cooperative symbiont, whereas when rewards are feedback-dependent (i.e., they are tied to the host benefit), population structure tends to favor symbiont cooperation. Furthermore, depending on the type of rewards and host benefits, both positive and negative frequency dependence might occur at different combinations of choosiness and population structure. Finally, population structure in a coevolutionary model of host choosiness and symbiont benefits tends to make the maintenance of cooperative symbionts and choosy hosts less likely. Overall, my results show that in contrast to its generally positive role in promoting cooperation in social evolution, population structure is not always good news for mutualisms with partner choice.

## 2 Model

I consider a well-mixed population of hosts that interacts with a subdivided population of two strains of symbionts that I will call Strains A and B. Each host individual encounters a local population in which Strain A is at frequency *q*, whereas the overall frequency of Strain A in the population is denoted by *p*. Population structure determines the distribution of this local frequency *q* given *p*, or *f*(*q|p*). For example, in a well-mixed population, *f*(*q|p*) would be binomially distributed. In structured populations, the distribution of *q* will be wider, so that a host will experience a wider range of local frequencies of the two strains. In particular, I assume that the symbiont population is patch structured, and with non-overlapping generations. This allows me to use well-known results from island-structured populations, in particular, the fact that the distribution of local frequencies of the two strains (corresponding to alleles) over patches is given by a beta distribution [44, 45, see equation (SI–5) in the Supplementary Information].

### 2.1 Partner choice

By convention, I assume Strain A is preferred by the host and also provides more benefit, so I refer to it as the “beneficial strain,” though both strains might be giving some benefits to the host. I assume that the host initially is infected by symbionts at their local frequencies ( *q* and 1 *− q* for Strains A and B, respectively; thus I ignore sampling variance by the host [46]). The host then “rejects” each infection by Strain B with probability *s*(0 *≤ s ≤*1), whereas infections from Strain A are always accepted. The variable *s* therefore measures the choosiness of the host. The new frequency of Strain A after partner choice, *q_c_*, is given by:

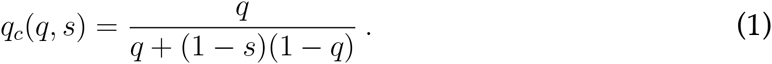

Figure 1 depicts the effect of partner choice in increasing the post-choice frequency of the beneficial symbiont.

**Figure 1:**
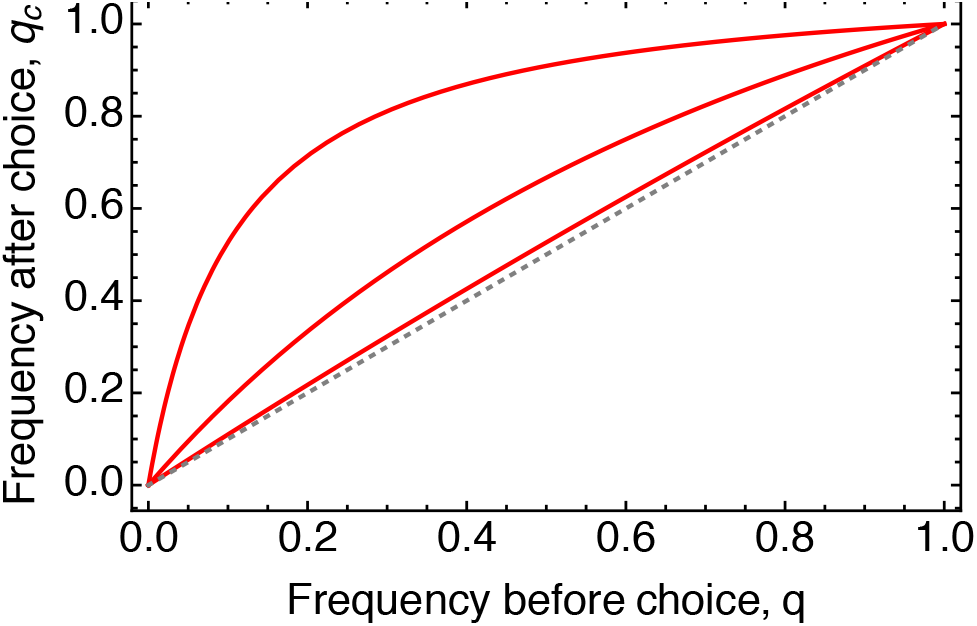
The frequency of Strain A (the “beneficial symbiont”) after choice, *q_c_*, as a function of its local frequency before choice, *q*. The dotted gray line is *q_c_*= *q*, whereas the solid red curves depict *qc*as given by equation (1) with (from top to bottom) *s*= 0.9,*s*= 0.5, and *s*= 0.1. Note that the enrichment of the beneficial symbiont (distance between the solid and dashed lines) is greatest at intermediate local frequencies.

### 2.2 Host fitness

I assume that the host fitness, denoted by *w_H_*, is a benefit *b*(*q_c_*) obtained as a function of *q_c_* minus some cost of partner choice, *c*(*s*):

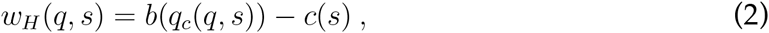

where I assume that *b*(*q_c_*) is an increasing function. The expected fitness of the host is then given by:

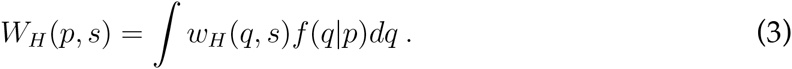

An approximation of this integral can be written by writing the host fitness in (3) in a Taylor series, and discarding terms of cubic order and higher:

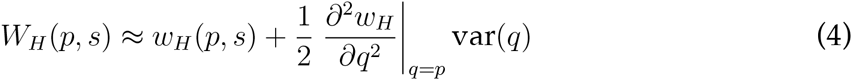

This approximation is particularly useful for our purposes because it decomposes the expected fitness into the fitness at the average symbiont frequency plus a term dependent on the variance of the local frequencies which is a measure of population structure: highly structured populations will have higher var(*q*), because local patches will be more likely to be enriched in one strain or the other. Thus, the coefficient of var(*q*), the second derivative of *w_H_* in *q* gives us the effects of increased population structure on the host fitness. This approximation is valid when the higher-order moments of *q* are small (i.e., when population structure is not too strong).

### 2.3 Symbiont fitness

I assume that symbiont fitness is derived from their interaction with the host through two potential reward mechanisms: fixed, and feedback-dependent. In fixed rewards, all symbionts that are not rejected by the host receive a fixed fitness increment from the host, denoted by *h_0_* ≥ 0, independent of the strain identity, the performance of the host or the symbiont frequency on the host. In contrast, the feedback-dependent reward results from the host providing rewards that are proportional to the benefits it receives, i.e. *h*_1_*b*(*q_c_*, where *h*_1_ ≥ 0 is a constant. I assume both strains experience the same rewards after the partner choice stage. Thus, a symbiont that survived partner choice gets benefit

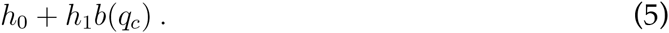

The beneficial strain, in consequence of providing more benefits to the host, incurs a cost *k* that the non-beneficial strain does not incur. Finally, I assume that symbionts that do not infect the host do not gain any rewards or pay costs. Thus, the per-infection fitness of the beneficial strain A in a local population with frequency *q* is:

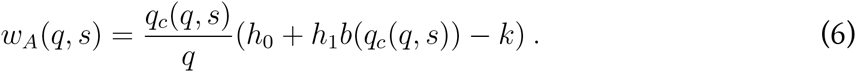

Likewise, for the non-beneficial strain B, we have:

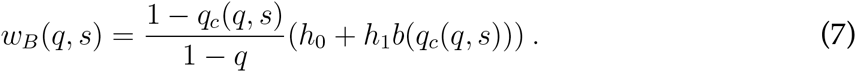

In (6) and (7), the fractions on the right-hand side account for the fact that partner choice enriches the beneficial strain on the host relative to the non-beneficial strain, and hence increases the relative per-capita fitness of the beneficial strain. The expected per capita fitness of the strains are given by:

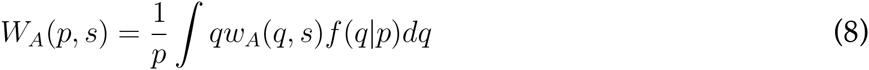

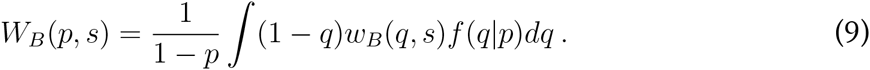

The sign of the difference *W_A_*(*p, s*) − *W_B_*(*p, s*) tells us whether symbiont A or B is favored by selection. Assuming selection is weak so that *f*(*q|p*) is given by a beta distribution [44, 45], the integrals (8) and (9) can be evaluated analytically, but the expressions are too cumbersome to reproduce here. For the presentation of results, I quantify population structure with the pairwise relatedness coefficient with non-overlapping generations [47, 48], given by:^1^

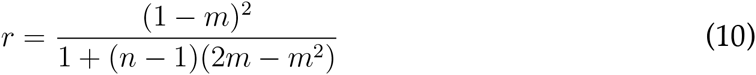

## 3 Effect of population structure on host fitness

As a baseline, I first compute the effect of population structure on a non-choosy host (with *s* = 0, i.e., always accepts all strains). As explained in section 2.2, the coefficient of the variance term in the Taylor expansion of expected host fitness gives the effect of population structure in the weak structure limit. In the absence of partner choice, this coefficient evaluates to:

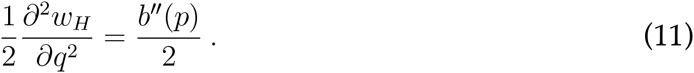

In other words, the variance term has the same sign as the second derivative of the host benefit function *b*(*p*). Therefore, stonger symbiont population structure will increase a non-choosy host’;s expected fitness if the benefit function *b*(*p*) is accelerating (*b″*(*p*) *>*0), and decrease if the benefit function is decelerating (*b″*(*p*) *<*0), in *q*. If host benefits are linear in *q*, the expected fitness of a non-choosy host is independent of population structure. When the host is choosy, i.e. *s* > 0, the coefficient of var(*q*) in equation (4) becomes:

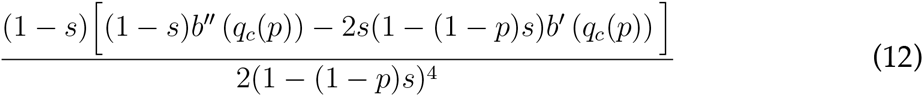

The denominator of this term is always positive, while in the numerator, the second term in the square brackets is always negative (since *b′*(*·*) *>*0 by assumption). Thus, the sign of the variance term in host fitness again depends on *b′′*(*·*), the shape of the host benefits as a function of *q*. With linear host fitness in *q*(b′′(*·*)=0), expression (12) is negative, meaning that host fitness decreases with population structure (increased var(*q*)). More generally, a sufficient condition for symbiont population structure to decrease host fitness is that *b*(*q*) is concave, i.e., the host’s benefit is a decelerating function of the frequency of the beneficial symbiont. Conversely, a necessary (but not sufficient) condition for population structure to increase a choosy host’s fitness that the benefit from the beneficial symbiont is accelerating (*b′′*(*·*) *>*0). These results indicate that partner choice tends to interact negatively with population structure in the expected host fitness, as the region where the host fitness increases with population structure is diminished with choice.

Figure 2 illustrates these results by evaluating the expected host fitness in equation (3) numerically for particular host benefit functions. It shows that, consistent with the analytical approximation, population structure tends to decrease the expected benefits to choosy hosts, except for low choosiness and accelerating benefits from the beneficial symbiont. This shows that for a given cost of choosiness, selection for host partner choice would be weaker or absent when symbiont populations are more strongly structured.

**Figure 2:**
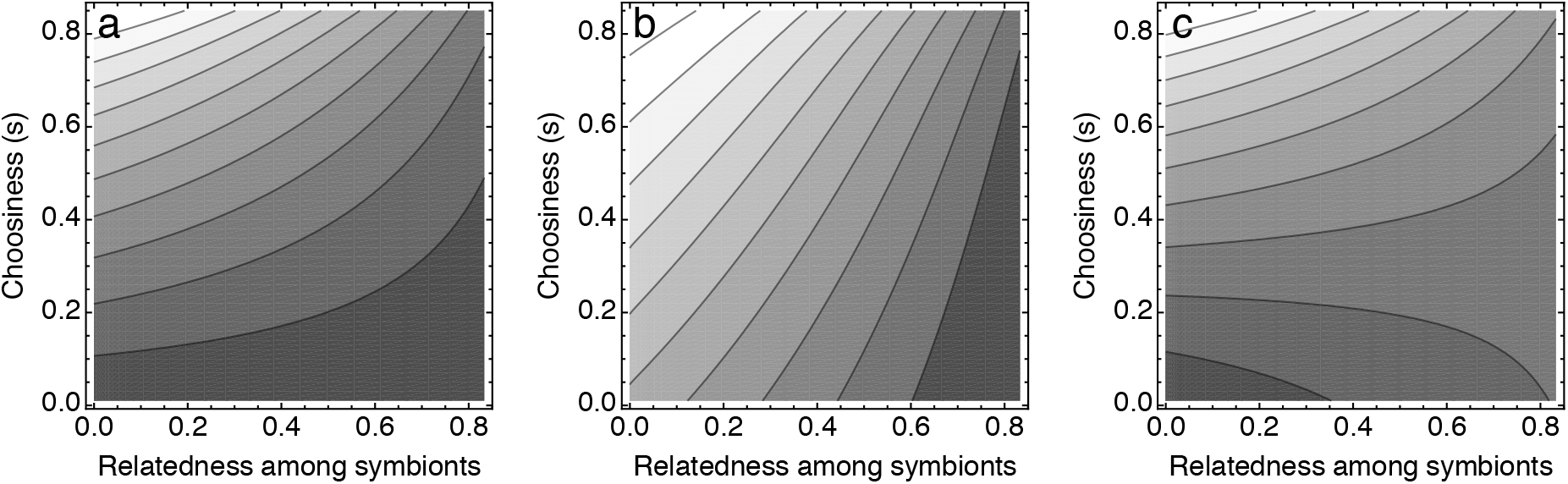
Increased population structure tends to decrease the expected benefit to choosy hosts. Here, I depict the host benefit as a function of host choosiness and the relatedness among symbionts as a measure of population structure, for three types of host benefit functions: linear (Panel a: *b*(*q_c_*)=*q_c_*, diminishing returns (Panel b: (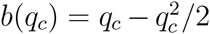)), and accelerating returns (Panel c: (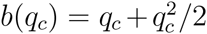)). Lighter shading indicates higher expected host benefit. For a given level of choosiness, increasing population structure (increasing relatedness) decreases host fitness when host benefits are linear or diminishing returns. For accelerating returns host benefits, increased population structure increases expected host benefit for low choosiness, but decreases for high choosiness. In all panels the patch size for symbionts, *n*= 100 and the overall frequency of the beneficial strain, *p*=0.5.

## 4 Effect of population structure on symbiont fitness

### 4.1 Fixed rewards: population structure hurts the beneficial symbiont

First, consider the case where the reward to each selected symbiont is fixed; in other words, I set the feedback coefficient *h_1_* = 0 in eq (5), while *h*_0_ > 0. I compute the expected fitness difference *W_A_*(*s*) − *W_b_*(*s*) using equations (8) and (9) at *p ≈*0 and *p ≈*1 to determine the invasion and fixation of the beneficial symbiont. Figure 3 shows the representative result: with increasing population structure, the beneficial symbiont becomes less likely to be able to invade and fix in a population of the non-beneficial symbiont. Figure 3 also shows that there exist combinations of choosiness and symbiont relatedness that create positive frequency dependence, where the beneficial symbiont cannot increase when rare but can go to fixation if it is common.

**Figure 3:**
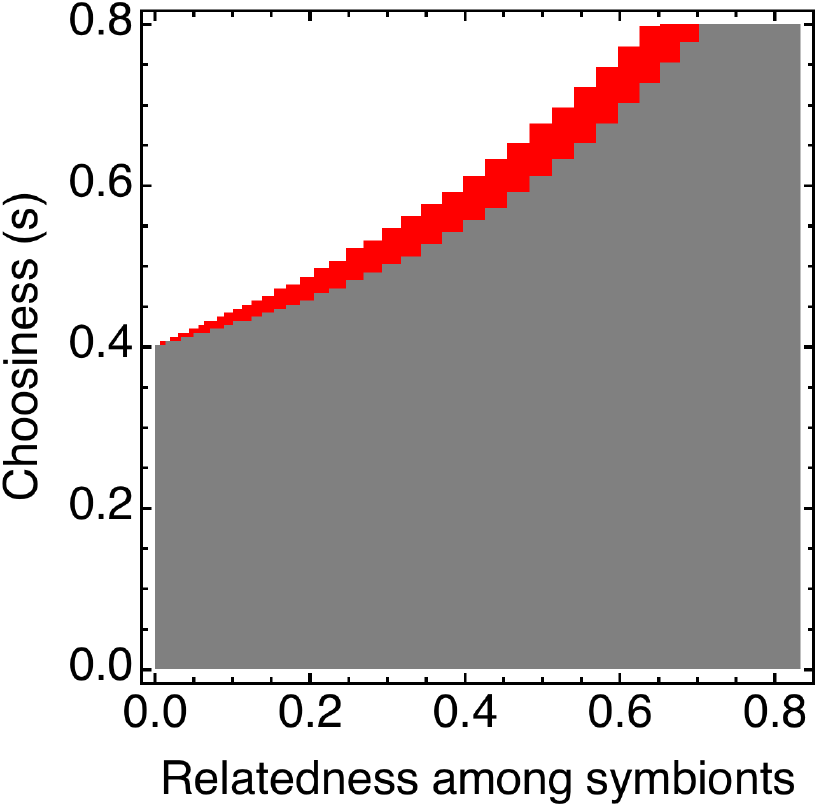
Incrasing population structure hurts the beneficial strain under fixed rewards from the host. The plot gives the combinations of host choosiness and relatedness between the symbionts that allow the beneficial symbiont to invade when rare and go to fixation when common in the population. Gray = beneficial symbiont can neither fix nor invade, red = beneficial symbiont can fix but cannot invade, white = beneficial symbiont can both invade and fix. For a given choosiness (going along a horizontal line), increasing relatedness between symbionts only makes the invasion and fixation conditions harder to satisfy. Parameter values are: *h_0_* = 1, *h_1_* = 0, *k* = 0.4, *N*= 100.

### 4.2 Feedback-dependent rewards: population structure helps the beneficial symbiont

Next, consider the case when the reward to the symbiont is entirely feedback-dependent, i.e., when *h_0_* = 0 while *h_1_* > 0. I consider two cases, first when the benefit to the host is a linear function of the beneficial symbiont frequency after partner choice, and second when the host fitness is a concave function, corresponding to diminishing returns from higher abundance of the beneficial symbiont. As above, I compute the expected fitness difference of the two strains using equations (8) and (9) at invasion and fixation of the beneficial strain. Figure 4 shows that unlike the fixed-rewards case, population structure with feedback-dependent rewards tends to favor the beneficial symbiont. In the linear host benefits case and without partner choice ( *s*= 0), the beneficial symbiont goes from being selected against at both invasion and fixation to selected for at a threshold relatedness value. With partner choice ( *s* > 0), the threshold relatedness for which the symbiont can both invade and fix goes down. Moreover, a region appears when the beneficial symbiont is favored when common but disfavored when rare, leading to positive frequency dependence.

**Figure 4:**
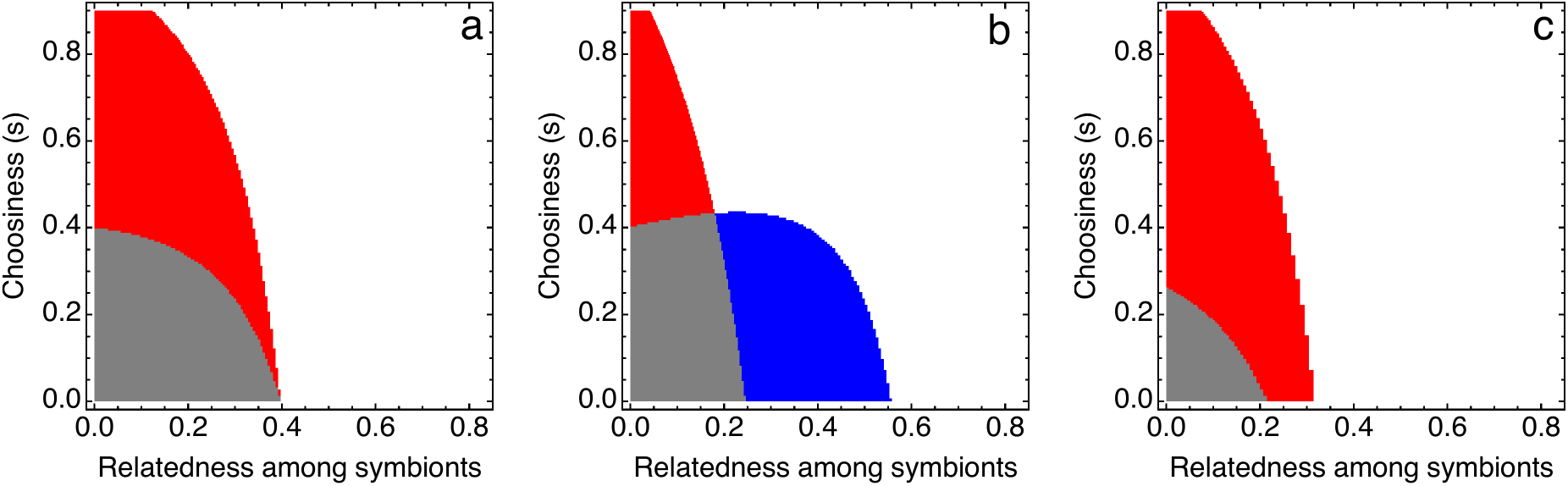
With feedback-dependent rewards, increasing population structure tends to help the beneficial symbiont. Panel a depicts the case where the host fitness is a linear function of the frequency of the beneficial symbiont after choice (specifically, *b*( *q_c_*) = *q_c_*, panel b a concave function (diminishing returns,(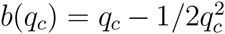)), and panel c a convex function (accelerating returns(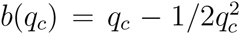)). Color scheme as in Figure 3: gray = beneficial symbiont can neither fix nor invade, red = beneficial symbiont can fix but cannot invade, white = beneficial symbiont can both invade and fix, blue = beneficial symbiont can invade but cannot fix. In all three panels, for a given choosiness (going along a horizontal line), increasing relatedness between symbionts tends to make the invasion and fixation conditions easier to satisfy. In all cases, high choosiness and low population structure tends to generate positive frequency dependence; with diminishing return benefits to the host, low choosiness and high relatedness can results in negative frequency dependency as well (in blue in the panel b). Parameter values for all panels are: *h_0_* = 0, *h_1_* = 1, *n* = 100. In panels a and c *k* = 0.4; in b, *k* = 0.2.

In the diminishing return case, a similar pattern obtains, with one major exception: at low choosiness, as relatedness increases the population transitions from one where the beneficial symbiont is selected against to one with negative frequency dependence where the beneficial symbiont is favored to invade but not fix (Figure 4b). At high choosiness, we again have a region of positive frequency dependence. At intermediate choosiness, there can also be a small region where increased relatedness hurts the beneficial symbiont, taking away the advantage it has when common.

Intuitively, these results can be understood by noting that the host benefit comes a public good through the rewards feeding back. In public goods dilemmas, increased relatedness tends to favor increased cooperation. Public goods with diminishing returns tends to produce negative frequency dependence, whereas those with accelerating returns generate positive dependence [51], consistent with our findings. Independently of this, as demonstrated by the fixed-rewards case, partner choice by itself tends to generate positive frequency dependence to the basic public goods selection patterns. Thus, depending on the combination of host choosiness, population structure, and host benefits, either kind of frequency dependence can obtain.

### 4.3 Local competition

The above analyses for selection on symbionts are based on the effect of the interaction on symbiont reproduction. However, it is well-known that factors such as local dispersal that create population structure can also create local competition between related individuals, which, depending on the mode of population regulation, can cancel out the effects of population structure on fertility [52, 53]. In the Supplementary Information SI–1, I consider the full life-cycle of the symbiont in the island model accounting for local competition. When local competition is taken into account, the effect of populations structure disappears with fixed rewards (due to the cancellation effect mentioned above). With feedback-dependent rewards, population structure hurts the beneficial symbiont at low to moderate host choosiness, but helps it at high host choosiness, regardless of the shape of the benefit curve.

## 5 Coevolution of host choosiness and symbiont cooperation

The above results are derived for the case where the host population is fixed for a level of choosiness. In this section, I consider the case where the host choosiness coevolves with symbiont strain frequencies. In particular, I again assume an island structure for both hosts and symbionts, and vary symbiont population structure while the host population stays well-mixed. I also assume a fixed cost to host choosiness (for a complete description of the model, see the Supplementary Information), and model the competition of a host genotype with choosiness *s* against a non-choosy host.

A population without the beneficial strain and non-choosy host is always a stable equi-librium. In addition to this equilibrium, the coevolutionary dynamics may also exhibit one of the following: a stable internal equilibrium where a polymorphisms of both the symbiont and host are maintained by selection, an unstable internal equilibrium, and a stable limit cycle.

A stable internal equilibrium obtains with both fixed and feedback-dependent symbiont rewards, when host fitness is a linear or concave (diminishing returns) function of the frequency of the beneficial symbiont (Figure 5, Supplementary Figure 2, 4). It requires that *s* is higher than a threshold (determined by the cost of symbiont cooperation) and the cost of host choosiness is not too high. The first ensures that for some frequency of choosy hosts beneficial symbionts can be maintained in the population, while the second for some frequency of the beneficial symbionts, choosy hosts can be maintained. The convergence to this internal equilibrium is always oscillatory. Even when a stable internal equilibrium exists, however, it is never globally stable: there always exists a region from which the population converges to the no-cooperation/no-choosiness equilibrium. A stable limit cycle can also exist with accelerating returns to the host and fixed rewards to the symbiont (not shown).

**Figure 5:**
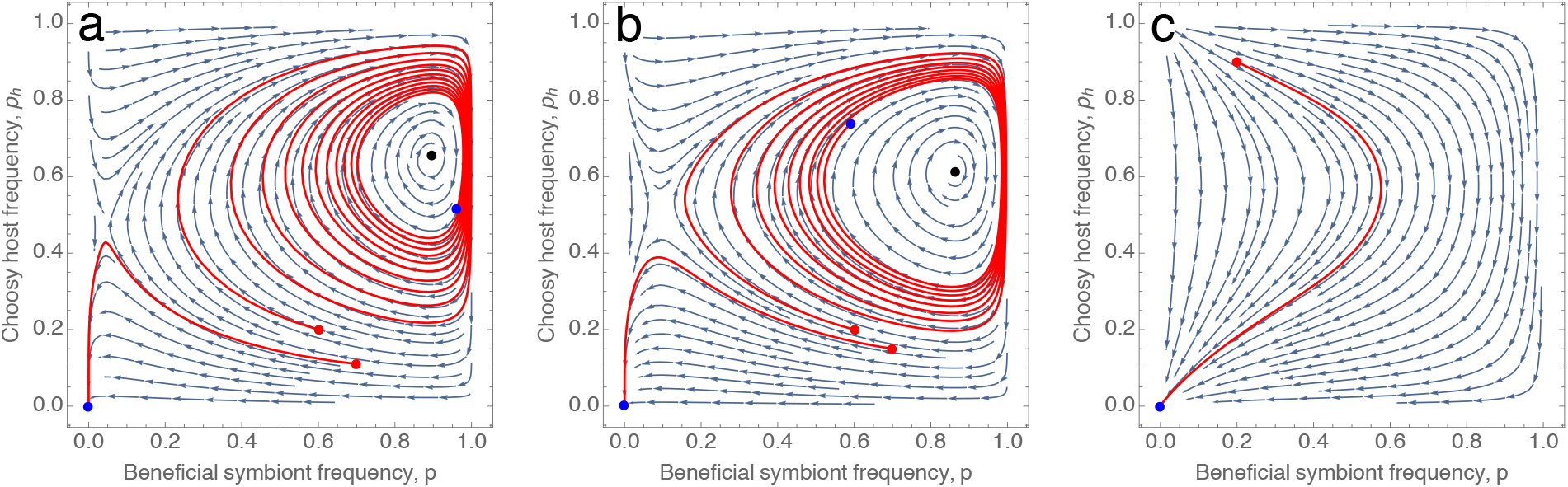
Coevolutionary dynamics of choosiness and symbiont cooperation show stable polymorphisms in both. The curved arrows depict the direction of frequency changes given by equations (SI–10) and (SI–14) while the red curves depict sample trajectories (starting at the red circles and ending at the blue ones). The black circles depict the location of the internal equilibrium. In all three panels, symbionts have fixed rewards and host benefit is linear (*b*(*q_c_*) = *q_c_*). In panel a, the symbiont population is well-mixed ( *m* = 1); panel b depicts the case with relatively strong population structure ( *m* = 0.01), while panel c is with very strong poulation structure ( *m* = 0.001). Population structure somewhat shrinks the basin of attraction of the stable polymorphism while also shifting it slightly downwards and left, before removing the internal equilibrium entirely; in other words, decreasing the expected cooperation maintained in the population. Parameters; *χ* = 0.1, *s* = 0.6, *h_0_* = 1, *h_1_* = 0, *b*(*q_c_*) = *q_c_*, *n_h_* = *n_s_* = 100.

The effect of population structure is to (i) decrease the stable frequency of the beneficial strain (Figure 6, though not for all parameter values for feedback-dependent rewards, see SI), (ii) to make the basin of attraction for this equilibrium smaller (Figure 5b), and (iii) when very strong, to make the internal equilibrium disappear entirely, leaving only the no-cooperation/no-choosiness equilibrium (Figure 5c).

**Figure 6:**
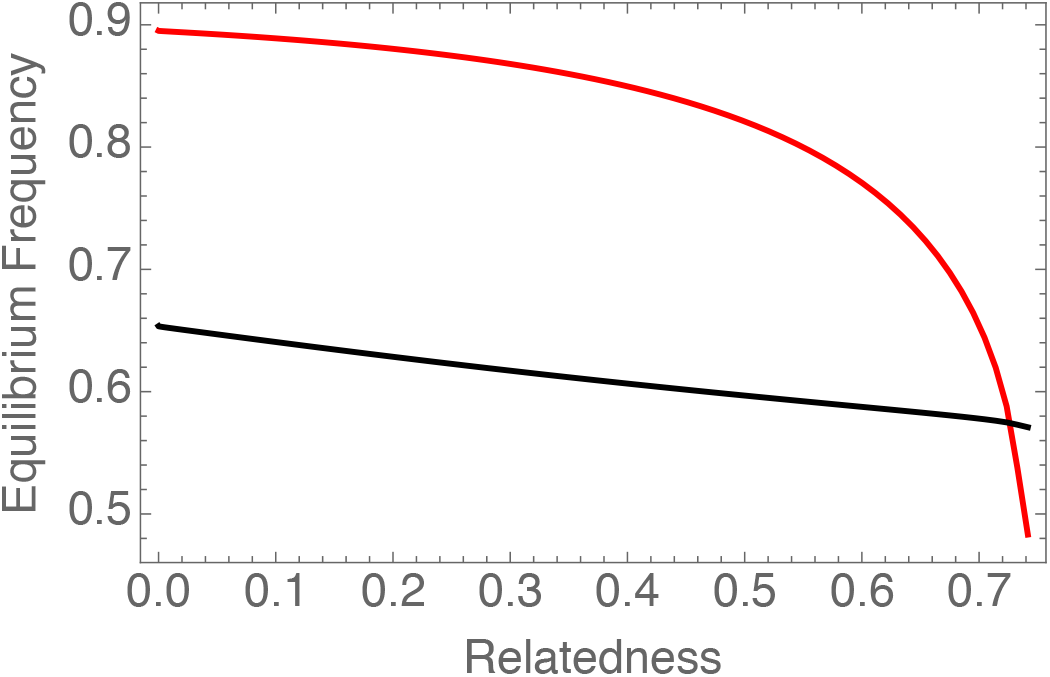
Population structure reduces the equilibrium frequencies of the beneficial symbiont and choosy host. This figure depicts the location of the stable equilibrium (red curve: symbiont frequency, black: host) for linear host benefits and fixed symbiont rewards (*h_0_* = 1, *h_1_* = 0). The equilibrium stays stable throughout, however, it disappears completely at very strong population structure. Other parameters as in Figure 5.

Overall, the coevolutionary model shows that depending on the type of benefit to the host and symbiont, polymorphisms of costly choosiness and symbiont cooperation can stably exist due to selection even without further (either through mutation or migration) input of variation into the population. At the same time, stronger population structure tends to make maintenance of cooperation and choosiness less likely.

## 6 Discussion

The basic question of this paper is how different mechanisms that generate fitness feedbacks interact with each other to promote or hinder the maintenance of cooperation in symbioses. In particular, I consider partner choice in conjunction with population structure and different kinds of rewards (fixed vs. feedback-dependent). My main results are the following: (i) for hosts, the benefit of partner choice is diminishing with increased population structure, except for fast enough accelerating benefits. For symbionts, (iia) when rewards from the interaction are fixed, population structure disfavors the beneficial strain. However, (iib) when rewards are feedback-dependent, population structure tends to generally favor the beneficial strain with potential for positive or negative density dependence, depending on the shape of the host benefit curve. (iic) Accounting for local competition removes the fixed reward effect of population structure, and while the beneficial effect with feedback-dependent rewards is conditional on high host choosiness. (iii) When host choosiness and symbiont cooperation coevolve, a stable polymorphism in both can be sustained by selection, but population structure tends to make this equilibrium and therefore the maintenance of cooperation less likely. Taken together, these results paint a complex pattern of interaction between these mechanisms commonly invoked to explain mutualism maintenance.

The main implication of my results for host evolution is that partner choice is less beneficial to hosts in more strongly structured populations. This effect follows directly from the logic of how partner choice works (that it requires local variation). Importantly, this effect is a direct one, i.e., independent of any changes in overall symbiont frequencies resulting from population structure. We do know that in many horizontally acquired symbioses, local pools of symbionts will vary between hosts due to population viscosity (as shown in, for example, rhizobia [54, 55] and mycorrhizae [56]). Such population viscosity is expected to drive local populations of symbionts to extreme frequencies, restricting the effectiveness of partner choice mechanisms. My coevolutionary results confirm this intuition: stronger partner choice tends to make less likely that equilibria where choice and beneficial symbionts are maintained are attained. This is because both the range of symbiont frequencies at which choice is worthwhile (for a given cost) diminishes and the threshold frequency of hosts required to favor the beneficial symbiont increases with increased population structure.

For the symbionts, my results reveal that population structure can hinder or promote selection for beneficial symbionts through partner choice, depending on what happens after choice. When each chosen symbiont gets a fixed fitness increment, population structure works against the beneficial strain. This scenario indicates that the interaction between partner choice and population structure by itself is negative: each reduces the effectiveness of the other. When feedbacks tied to the host’s performance (sometimes termed partner fidelity feedbacks) are added to the mix, population structure can promote selection for the beneficial strain. This is consistent with the fact that the feedback rewards from the host are effectively public goods for the symbiont, and increased relatedness favors production of public goods [41, 42].

The intuition that population structure might reduce local variation available to partner choice has been pointed out by Verbruggen et al. [17], but they were concerned explicitly with within-host genetic structure of the symbiont, as opposed to population-level structure that I focus on. Another difference between the current work and Verbruggen et al. lies in the fact that I derive the fitness consequences of partner choice in structured population from a simple but generally applicable model of partner choice in structured population, rather than assuming functional forms of relative fitness, as Verbruggen et al. do. These differences notwithstanding, my model is closest to the “coarse control” model of Verbruggen et al., because I assume that hosts (after choice) reward each symbiont partner equally (whether through fixed- or feedback-dependent rewards). It would be interesting to see how my results might change when hosts can provide directed or negotiated [29, 57] rewards to only beneficial symbionts after choice. In both cases, there is potential for negative interaction between population structure and directed rewards after choice [29, 42].

Kiers et al. [58] discuss the importance of spatial structure in maintaining cooperation in the agriculturally (in addition to ecologically) important plant-soil symbioses. They mainly focus on the conflicting effects of relatedness and local competition in promoting and opposing cooperation, indicating that the overall effects of agricultural practices like tilling that reduce population structure can be theoretically ambiguous. My results (as depicted in Figure 2) suggest that increased population structure (e.g., a no-till agricultural strategy) will directly reduce average plant performance (before any changes in symbiont frequencies), but if symbiont fitness is mainly driven by feedback-dependent rewards, this direct effect can be offset by an increase in the frequency of beneficial strains. The evidence reviewed by Kiers et al. mostly indicates higher mutualistic benefits from more structure oil populations, but no direct study exist to my knowledge that attempts to tease apart the direct effects of population structure from the effects of subsequent symbiont evolution.

My results also have implications for how variation in mutualisms can be maintained in the face of mechanisms that favor more cooperative partners. This problem has recently come to the forefront of theoretical and empirical research [37, 39, 40, 46]. My results provide three qualitatively different scenarios in which variation in the symbiont can be maintained. First, under feedback-dependent, but diminishing rewards from the host, which in accordance with public goods games theory [51], tends to generate negative frequency dependence (Figure 4, right-hand panel). Second, with fixed or feedback-dependent and linear or accelerating rewards, where we observe positive frequency dependence, and therefore bistability (Figures 3,4 [left-hand panel], and 1 [right-hand panel]). Such bistability can lead to a geographic mosaic of coexistence over larger spatial scales [59, 60]. Third, when host choosiness is coevolving at the same time-scale as symbiont frequencies, there can be stable polymorphisms in both hosts and symbionts. Generally speaking, stronger population structure tends to make all three scenarios less likely by favoring one or the other strain unconditionally, or by removing polymorphic coevolutionary equilibria.

Finally, the models presented in this paper leave some questions open that will be useful to explore in future research. The first is that, for simplicity, I did not consider the joint spatial structure of hosts and symbionts. The build up of genetic correlations between partners is likely to have an impact on the conclusions about the effects of population structure [24], although preliminary simulations (not shown) suggest that the qualitative patterns might be similar. Likewise, I did not consider the potential for post-infection partner choice or rewards directed specifically to beneficial symbionts. Such directed rewards, depending on how they are determined (e.g., absolute vs. relative [42] or negotiated rewards [29]) can reinforce or counteract the effects of partner choice under population structure.

In conclusion, my results show that multiple mechanisms of fitness feedbacks can interact in complex ways in selecting for or against cooperation between species and variation thereof. Understanding these interactions will lead to a more complete picture of the evolutionary and ecological dynamics of mutualisms.

## Acknowledgements

I thank E. Simms and J. Van Cleve for valuable comments on the manuscript.

## Supplementary Information

### SI–1 Full life-cycle in the island model

Here, I derive the expression for the change in frequency of the beneficial symbiont in an island model with weak selection. Consider a deme that contains the beneficial symbiont at frequency *q*. The fitness of the beneficial symbiont is given by:

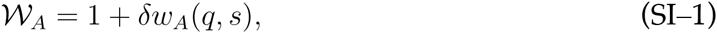

and for the non-beneficial symbiont:

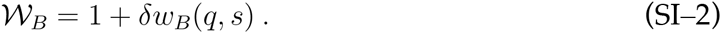

The new local frequency of the beneficial symbiont, after selection and migration is:

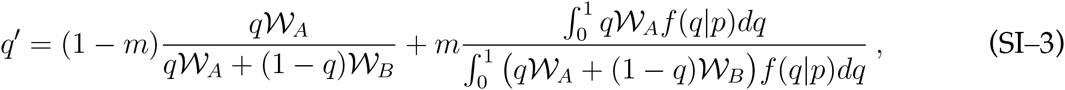

where *m* the the probability of new individuals in a patch being migrants. Taking the Taylor expansion to first order in *δ*, and integrating over *f*(*q|p*), we find the new overall frequency of the beneficial symbiont, *p′*:

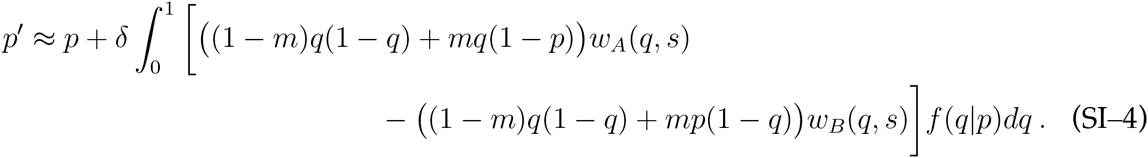

Finally, I assume selection is weak enough that the distribution of the local deme frequencies of the beneficial symbiont, *f*(*q|p*), is well-approximated by neutrality [1]. Thus, we have:

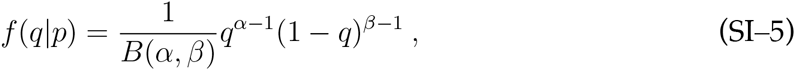

where *B*(*·, ·*) is the Beta function, *α*=*nmp*, *β*=*nm*( *p −*1), and *n*is the symbiont deme size.

Local competition changes the results reported in the main text section 4.1 and 4.2 in substantive ways. Under fixed rewards, population structure has no influence on whether selection favors the cooperative symbiont or not: there exist a threshold level of choosiness independent of population structure above which the beneficial symbiont is favored and below which it is not (Supplementary Figure 1, left-hand panel). This is a case of fertility effects of population structure being cancelled out exactly by the local competition effect [2]. On the other hand, under feedback-dependent rewards, population structure helps the beneficial symbiont at high levels of choosiness while hurting it at low levels (Supplementary Figure 1, right-hand panel). In particular, there exists a (low to moderate) range of host choosiness where the beneficial symbiont goes from being favored only when common to never being favored as population structure increases, whereas at higher host choosiness levels the symbiont goes to being always favored. Interestingly, and in contrast to the results without local competition, this pattern holds regardless of the shape of the host benefit. The reason local competition does not exactly cancel out the fertility effects under feedback-dependent but not fixed rewards is that in the former, there are effectively two social interactions involved: the first is partner choice, while the second is a public goods game through the host. Due to partner choice, the second stage happens over a modified distribution of strain frequencies relative to the distribution over which the local competition happens, therefore the the latter does not entirely cancel the former: when choosiness is low, local competition becomes more important as population structure increases, whereas when choosiness is high enough, the benefits from the public goods rewards become more important.

### SI–2 Coevolutionary model

In this section, I describe the model where two genotypes each of the host and symbiont exist in the population. One of the host genotypes has choosiness *s* and the other zero (i.e, does not reject any symbiont). I assume that the choosy host’s fitness when the local frequency of the beneficial symbiont is *q*is given by

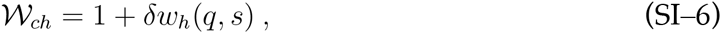

where *δ* is the strength of selection, assumed to be small, *w_h_* is given by equation (2), and I assume a linear cost to choosiness:

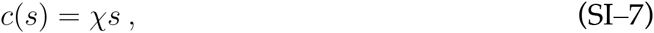

with *χ* a constant. Because of the weak selection assumption, I use the neutral distribution of local frequencies for both the host and symbiont [e.g. 1, 3]. In particular, I assume that the host population is well-mixed. I denote the distribution of the local frequencies of the choosy host by *g*(*q_h_*|*p_h_*). This assumption that hosts are well-mixed among patches allows me to neglect potential build up of correlations between host and symbiont genotypes [4] so that the joint local frequency distributions of hosts and symbionts are given just by the product of their separate distributions. The fitness of a non-choosy host is:

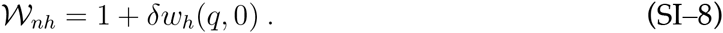

The overall frequency of choosy hosts after selection is

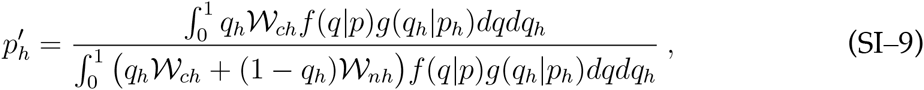

which to first order in the strength of selection becomes:

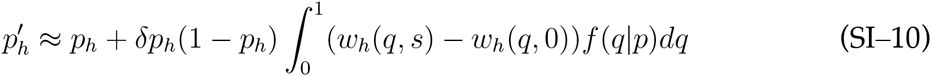

Now I turn to the symbiont fitness. In a deme that contains the beneficial symbiont at frequency *q* and the choosy host at *q_h_*, the fitness of the beneficial symbiont is given by:

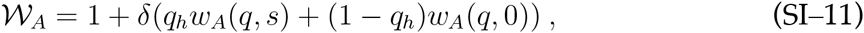

and for the non-beneficial symbiont:

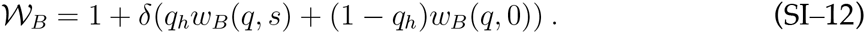

The new frequency of the beneficial symbiont, after selection and migration is:

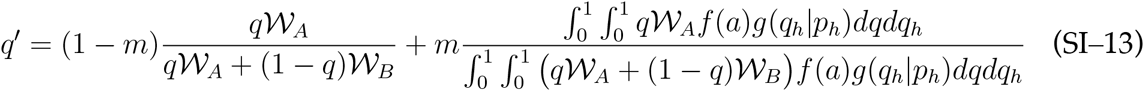

Taking the Taylor expansion to first order in *δ*, and integrating over *f*(*q|p*)and *g*(*q_h_*δ*p_h_*), we find the new overall frequency of the beneficial symbiont, *p′*:

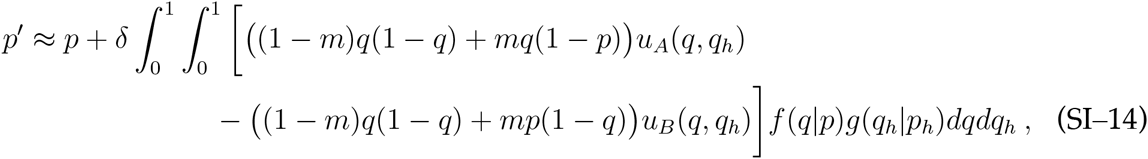

where *u_A_*(*q, q_h_*) = *q_h_w_A_*(*q, s*)+(1– *q_h_*)*w_A_*(*q* 0) and *u_b_*(*q, q_h_*) = *q_h_w_B_*(*q, s*)+(1–*q_h_*)*w_B_*(*q*, 0), i.e, the expected fitness increments to each strain due to the interactions in the local deme. The local choosy host frequency distribution is given by:

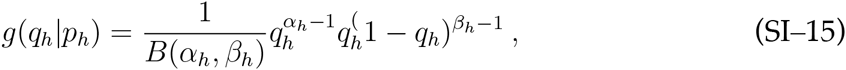

where *α_h_* = *n_h_p_h_*, *β_h_* = *n_h_*(*p_h_*–1), *n_h_* are the symbiont and host deme sizes, respectively[1].The symbiont local frequency distribution *f*(*q|p*) is again given by (SI–5).

Below, I sketch out the general patterns of the coevolutionary dynamics of the choosy host and beneficial symbiont frequencies, starting with fixed rewards to the symbiont.

**Supplementary Figure 1:**
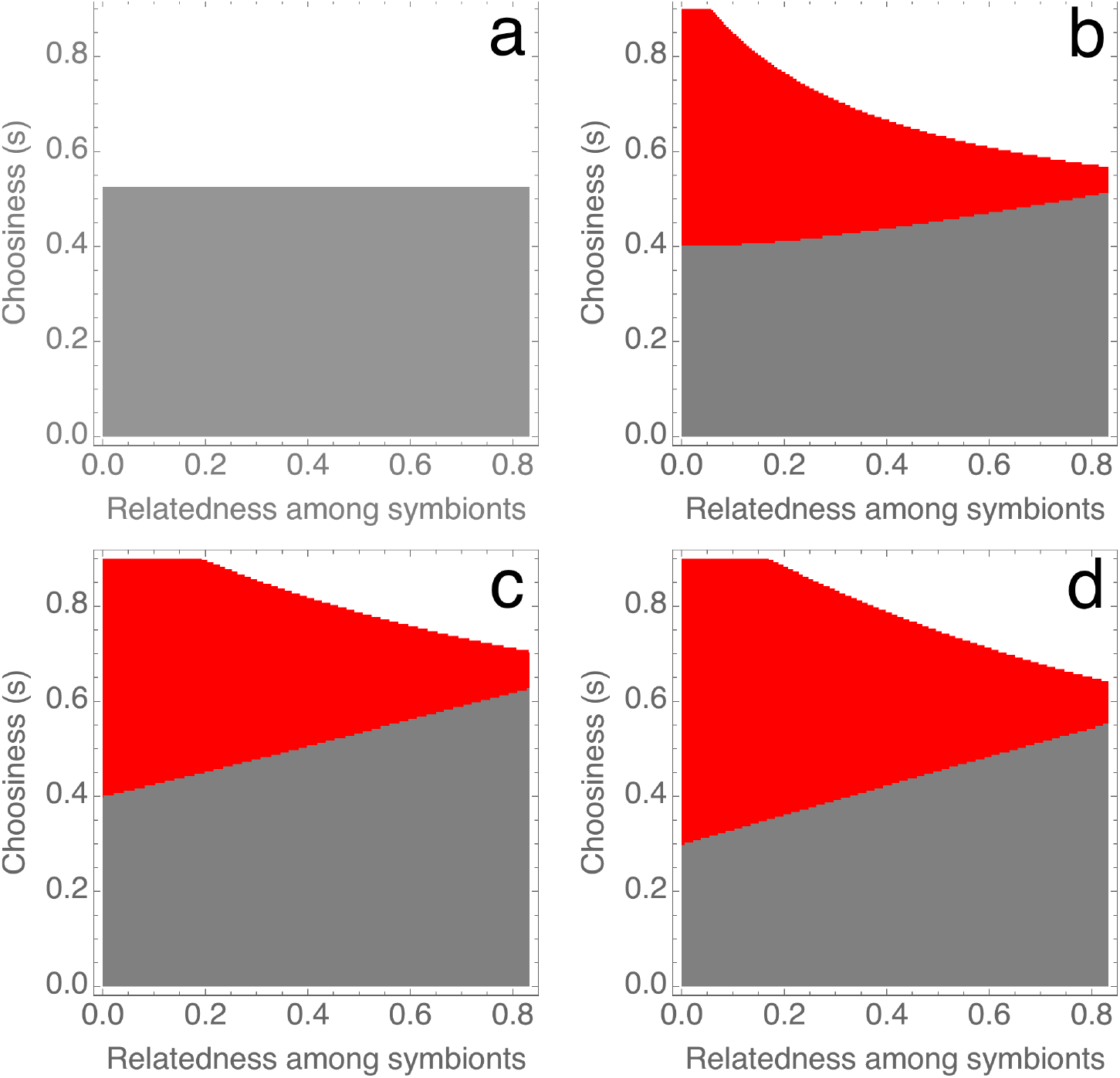
Effect of population structure changes when local competition is taken into account. Colors as in Figures 3 and 4 in the main text (gray=beneficial symbiont does not invade or fix, white=beneficial symbionts invades and fixes, red=beneficial symbiont cannot invade but can fix). Panel a depicts the fixed rewards (*h_0_* = 1, *h_1_* = 0) case: population structure has no effect on selection in the symbiont population. Other panels depict the feedback-dependent reward cases (*h_0_* = 0, *h_1_* = 1) with different shapes of host benefit. The all show that with high host choosiness, population structure helps the beneficial symbiont getting selected for at low frequencies while for lower host choosiness, the symbiont goes from being favored when common to not being favored ay any frequency. Particular host benefit functions are as follows. Panel b: diminishing returns (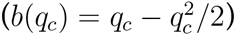), c: linear returns (*b*(*q_c_*)=*q_c_*), d: accelerating returns (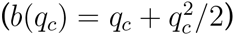). In all panels *k* = 0.4 and *n* = 100, except for the diminishing rewards (panel b), where *k* = 0.2.

**Supplementary Figure 2:**
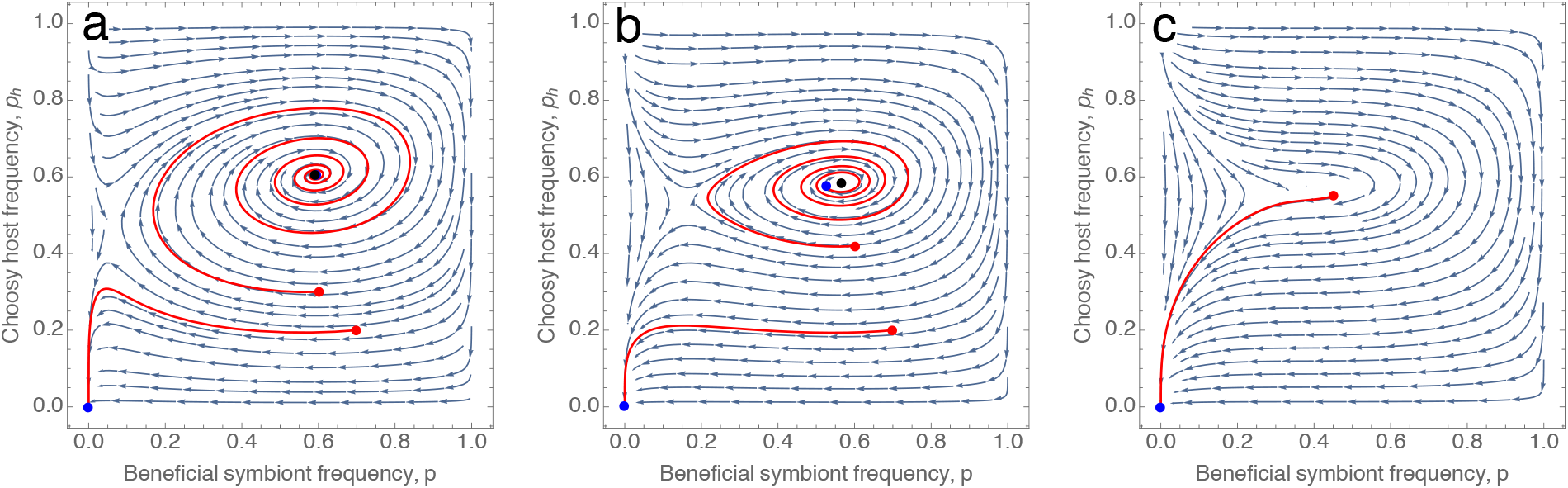
Coevolutionary dynamics with fixed benefits to the symbiont and diminishing returns to the host. Parameters for the three panels same as in Figure 5 in the main text, except 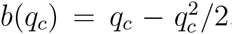. As in the linear host benefit case, population structure reduces the basin of attraction of the stable equilibrium before destroying it completely.

**Supplementary Figure 3:**
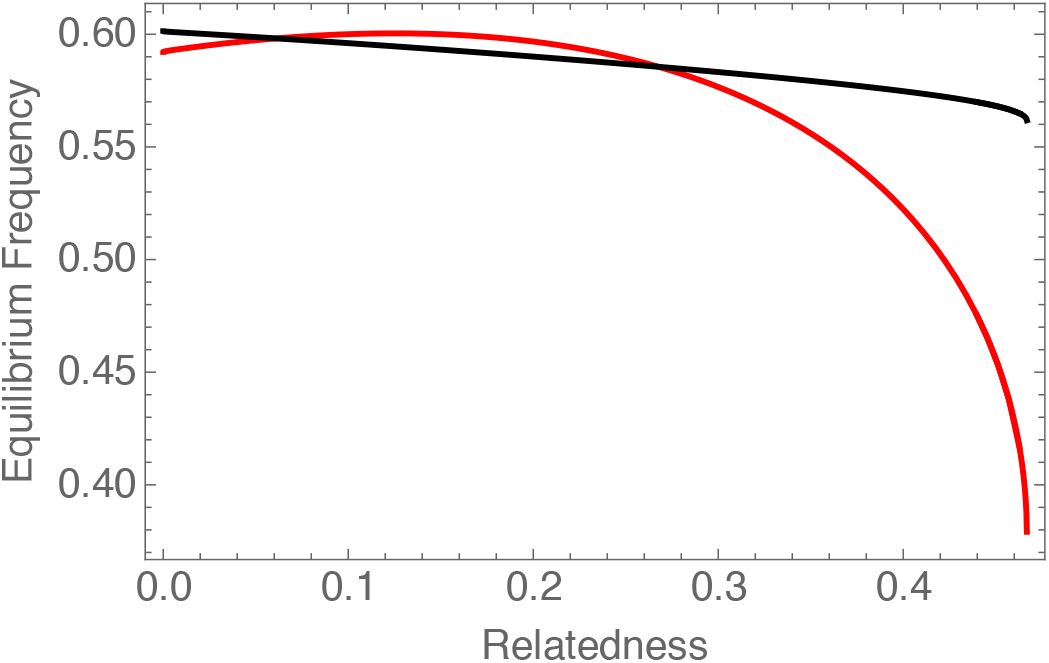
With diminishing returns to the host and fixed rewards to the symbiont, population structure might first increase, then decrease the frequency of the beneficial symbiont. Red curve gives the equilibrium frequency of the beneficial symbiont; black the choosy host. Parameters as in Supplementary Figure 2.

**Supplementary Figure 4:**
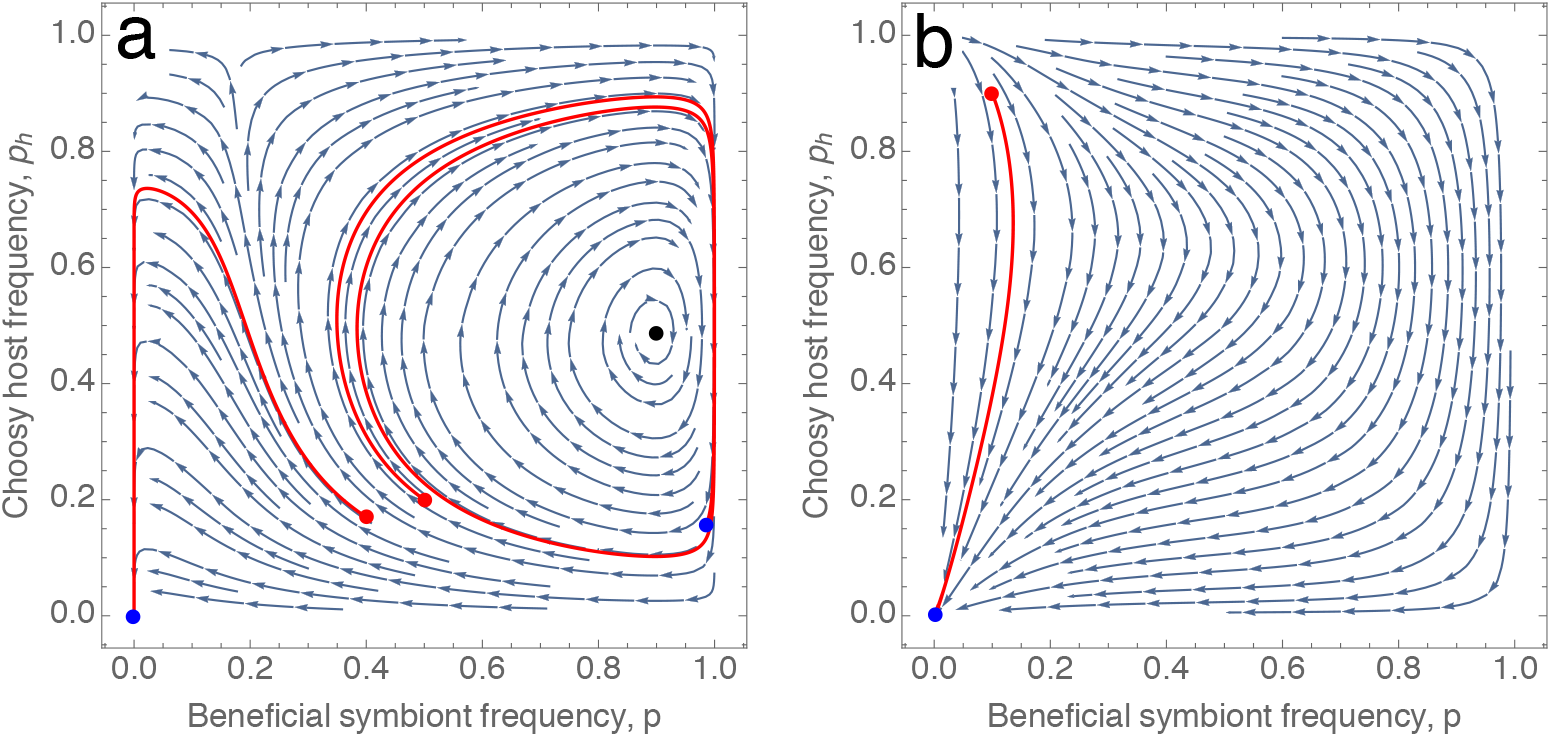
Population structure can eliminate the stable internal equilibrium with feedback-dependent rewards. The filled black circle in panel a depicts the stable equilibrium, which disappears in panel b. This figure assumes linear benefits to the host (*b*(*q_c_*)=*q_c_*) and feedback-dependent symbiont rewards (*h_0_* = 0, *h_1_* = 1). Host choosiness is *s* = 0.8; *m* = 1 on the left-hand panel; *m* = 0.001 on the right-hand. Other parameters: *χ* = 0.1, *k* = 0.4, *n_h_* = *n_s_* = 100.

**Supplementary Figure 5:**
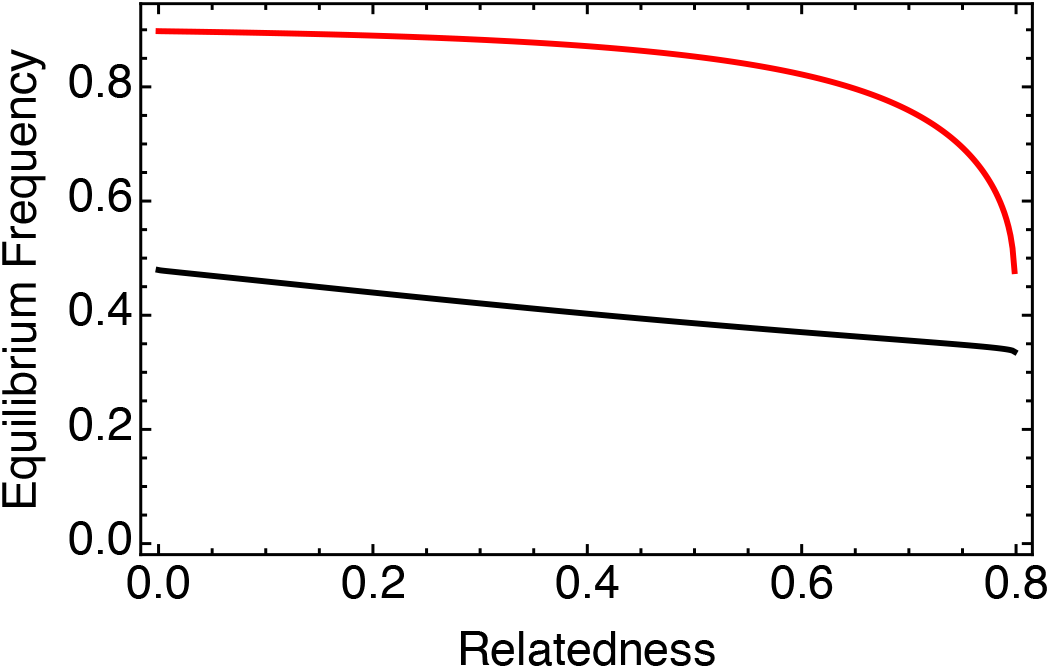
Population structure decreases the equilibrium frequency of the beneficial symbiont and the choosy host with feedback dependent rewards. This figure assumes linear benefits to the host (*b*(*q_c_*) = *q_c_*) and feedback-dependent symbiont rewards (*h_0_* = 0, *h*_1_ = 1). Other parameters as in Supplementary Figure 4.

#### 1 Fixed rewards to the symbiont

The first finding for fixed rewards to the symbionts is that there exists parameter combinations for which polymorphisms in both host and the symbiont are stably maintained, even in the absence of mutational (or migrational) input which most previous research has suggested to be important. However, the equilibrium where the beneficial symbiont and choosy hosts go extinct also is always stable. Stronger population structure of symbionts shrinks the basin of attraction of the stable equilibrium and shifts the equilibrium frequencies of the beneficial symbiont and choosy hosts lower, as shown in Figure 5 in the main text, which depicts the case where host fitness is a linear function of *q_c_*, and Figure 6, also in the main text, which depicts how the equilibrium frequency changes with population structure.

This pattern stays largely the same when host benefit exhibits diminishing or accelerating returns to *q_c_*(Supplementary Figure 2), although with diminishing returns, population structure can initially increase the equilibrium frequency of the beneficial symbiont (Supplementary Figure 3). Accelerating returns tend to make the cycles larger for the same parameter set.

To summarize, the overall effect of population structure in the fixed rewards case is to reduce the amount and likelihood of stable cooperation. The model also predicts that the coexistence patterns will generally exhibit at least transitory cycling when starting in the interior, either as oscillatory convergence to the stable equilibrium or as limit cycles. This is in line with previous results [5, 6].

#### 2 Feedback-dependent rewards

The pattern with feedback-dependent rewards is broadly similar to fixed rewards. The boundary equilibrium without the beneficial symbiont and choosy host is again always stable. There can also be a stable internal equilibrium as in the fixed-rewards case, but for the same parameters (except for *h_0_* and *h_1_*) a higher level of host choosiness is required for this equilibrium to exist. Population structure has the same effects as in the fixed-rewards case: it shrinks the basin of attraction of the stable equilibrium, eventually removing the stable equilibrium (Supplementary Figure 4), while also generally tending to reduce the frequency of cooperation and choosiness (Supplementary Figure 5).

## Footnotes

1 Note that I use this relatedness coefficient not to calculate fitness effects as is commonly done in social evolution theory, but simply as a familiar measure of population structure. The expectation over the distribution of deme frequencies *f*(*q|p*) accounts for the inclusive fitness effects [49, 50].

